# Prognostic significance of Substance P/ Neurokinin 1 receptor and its association with hormonal receptors in breast carcinoma

**DOI:** 10.1101/2020.06.27.175083

**Authors:** Riffat Mehboob, Syed Amir Gilani, Amber Hassan, Imrana Tanvir, Rizwan Ullah Khan, Shaista Javaid, Javed Akram, Sadaf, Fridoon Jawad Ahmad, Miguel Munoz

## Abstract

To evaluate the expression and Immunolocalization of Substance P (SP)/ Neurokinin-1 Receptor (NK-1R) in Breast Carcinoma (BC) patients and it’s association with routine proliferative markers (ER, PR, HER2/ neu and Ki-67).

**Methods:** A cross-sectional study was performed on 34 cases of BC. There were 23 cases of group A (Grade III), 8 of group B (Grade II) and only 3 cases of group C (Grade I). Age range comprised of patients from 20-80 years and the mean age of patients was 45.74 years. HE, ER, PR, HER2 and Ki-67 staining was performed as routine biomarkers. Samples were then processed for immunomarkers study of Substance P and NK-1R immunohistochemistry was performed for few cases.

**Results:** 14/23 cases (61%) of group A, 7/8 cases (88%) of group B and 2/3 (67%) cases of group C were SP positive. Overall, strong staining (≥ 10% tumors cells), labeled as “3+”, was observed in 9/14 (64.2%) cases of group A and 1/8 (12.5%) case of group B. Moderate staining labelled as “2+” (in ≥ 10% tumor cells) was observed in 3/14 (21.4%) cases of group A, 4/8 (50%) cases of group B. weak positive staining “1+” was observed in only 2/14 (14.28%) cases of group A, 2/8 (25%) cases of group B and all 2/2 (100%) cases of group C.

**Conclusions:** SP and NK-1R is overexpressed in breast carcinomas and there is significant association between grade of tumor and their overexpression. It may serve as a novel biomarker for grading of BC. We also suggest that NK-1R antagonists as a potential therapeutic strategy to inhibit and manage BC.

**Key Points:**

- Immunohistochemical expression of Substance P and Neurokinin 1 Receptor in breast carcinoma tissue was evaluated
- It was strongly expressed in grade III, with maximum intensity
- It may be investigated further for its role as prognostic and diagnostic marker
- Therapeutic potential of Neurokinin-1 Receptor antagonists must be explored

## INTRODUCTION

Breast cancer (BC) is most common cancer in women all over the world with an incidence of approximately 2 million in 2018. The highest rate of BC was observed in Belgium with 113.2/ 100, 000 women [1]. It can occur as a result of cells under the influence of estrogen multiplying and infringing on other tissue spreading to other regions of the body [2]. Invasive lobular carcinoma is the second most common type of BC, several histological sub types exist, most of the tumors are classified as grade II and majority of grade III are among the non-classified subtypes showing disease free region as compared to grade II [3]. The number of positive axillary lymph nodes and hormone receptor negative tumors increase among grade III tumors[4].

In BC, the malignant cells are enlarged with vacuolated cytoplasm and vesicular nuclei containing prominent nuclei. Most of the time the stroma was found to be increased and degenerative in nature[5]. There are various types of BC; they are classified as in situ and invasive. In situ carcinoma includes lobular carcinoma in situ (LCIS) and ductal carcinoma in situ (DCIS). Invasive carcinoma includes Invasive Lobular Carcinoma (ILC) and Invasive Ductal Carcinoma (IDC) [6]. The grading of invasive BCs is the important factors in addition to size and the status of the lymph nodes [7]. Benign breast diseases especially fibroadenomas are also important as some of them (30%) may lead to cancer [8].

The staging of BC is related to the size, location and number of regional metastases to lymph nodes and sometimes are related to growth [9]. TNM Stage IIB, IIIA, and IIIB are tumor stages help in diagnosis [10]. BC is commonly caused by low-penetrance genes that are involved in DNA repairing mechanism. DNA damage and chromosomal damage may also cause BC The XRCC3Thr24Metpolymorphism is the most common gene associated to BC [11]. These are repair genes to rectify the DNA damages. These genes are involved in enhancing the cytotoxicity, apoptosis, p53 phosphorylation and exposure to external factors that cause DNA damage [12]. In stage 2 about 54% of the women are diagnosed, while in stage 1 only 16% are diagnosed [13]. Worldwide the occurrence of BC exceeds all female cancers with high mortality rates [14].

Substance P (SP) is a small undecapeptide hormone [15] and most abundant tachykinin (TK) peptide in the central nervous system of mammals [16]. Many physiological and pathological roles of this peptide have been noticed [17]. Munoz and Covenas [18] suggested a strong role of SP-Neurokinin-1 Receptor (NK-1R) system in the progression of carcinogenesis. BC cells exhibit mRNA for the receptor of SP, NK-1, which is then involved in promoting the cell proliferation and consequently metastasis [19]. Additionally, SP is also involved in vasculogenesis, angiogenesis and neo-angiogenesis as observed in both *in vivo* and *in vitro* studies, an essential step towards invasion and metastasis [20, 21].

It is first study to report the expression and distribution of SP in BC and to suggest a strong association of its expression with the progression of disease and its association to routine proliferative and hormonal markers. Thus, the aim of this study is to evaluate the expression of SP/NK-1R and its relationship with tumor type and clinicopathological parameters of breast cancer patients. Furthermore, the relationship between the SP/NK-1R and proliferative markers were investigated.

## Material and Methods

We have followed the same methods for data collection and immunohistochemistry as done in our previous study[22]. Study setting was Faculty of Allied Health Sciences, The University of Lahore, Lahore, Pakistan. A total of 34 formalin fixed paraffin embedded (FFPE) blocks of BC were included. Medical and personal history of patients consisted of age, span of disease, tumor site/size, progression of disease, staging/grading etc. Age range was 20-80 years. For collection of data, we followed American Joint Committee for cancer staging and End results reporting. All the parameters of Declaration of Helsinki were respected in this study. Classification of tumor was based on WHO criteria such as Well differentiated, moderately differentiated and poorly differentiated breast carcinoma for grade I, grade II and grade III respectively. All the slides were routinely stained with Hematoxylin-Eosin to assess the morphology of cells and proper classification of cases. These were interpreted by two histopathologists.

### ER, PR, HER2 and Ki-67staining

Immunohistochemistry (IHC) for PR, ER, HER2/neu and Ki-67 was accomplished on FFPE tissue segments as part of the routine clinical assessment of these cases using anti-ER antibody (DAKO, Denmark),anti-ER antibody (DAKO, Denmark), anti-HER2 (1:400 to 1:600, DAKO, Denmark), anti-PR antibody (DAKO, Denmark) and Ki-67 (DAKO, Denmark) with Visualize system for detection. Lobular and ductal normal areas of breast were as control for ER, PR and HER2 IHC whereas appendix tissue was set as control for Ki-67. Olympus (Model U-DO3) was used for microscopy.

### Substance P/NK-1R Immunohistochemistry (IHC)

FFPE sections of 4μm were deparrafinized with xylene and decreasing grades of alcohol, washed in distilled water and then Phosphate Buffer Saline (PBS). These sections were pretreated with citrate buffer in microwave and were allowed to cool for atleast 20 minutes. Washings in distilled water and PBS was done before 3% H_2_O_2_ (30 minutes) to block the endogenous peroxidase activity. SP antibody (Biogenex) in dilution 1: 100 and NK-1R antibody (Abcam) in 1: 100 dilution was applied to the sections for 45-50 minutes in humid chamber. Washing step in PBS was done for 10-15 min. Slides were then incubated with secondary antibody Horse Raddish Peroxidase (HRP) for 45-50 minutes and washed again with PBS (10-15 minutes). 3,3’-Diaminobenzidine (DAB) is applied for 5-10 minutes and counter stained with hematoxylene for 2 minutes. FFPE sections were dipped in increasing grades of alcohol and then xylene for 5 minutes each. DPX mounting medium was used and slides were cover slipped. Methods are similar to one of our previous study on oral squamous cell carcinoma[22].

### Grading of IHC

Cell counting at 10 and 40X was done for the evaluation of protein expression and counts were made as in our previous study (table 1)[23]. Scoring for ER, PR, HER2 and Ki-67 was done by ALLRED method proposed by Qureshi and Pervez [24] (Table 2). No protein overexpression or membrance staining in < 10% tumor cells were labeled as score “0” and considered negative for SP/ NK-1R protein overexpression. Faint/weak staining (in ≥ 10% of tumors cells) were given the “1+” score.

**Table 1:** Interpretation of ER, PR and HER2 by Allred method

| ALLRED | Cell stain |  | Proportion score |
| --- | --- | --- | --- |
|  | % | Score (3) |  |
| Negative | 0 | 0 | 0 |
| Weak positive | 1 | 1 | 1 |
| Moderate positive | 1-10 | 2 | 2 |
| Strong positive | 10-33 | 3 | 3 |
|  | 33-66 |  | 4 |
|  | 66-100 |  | 5 |
| **Sum of proportion score and intensity score** |  |  |  |
| Negative |  | 0-2 |  |
| Positive |  | 3-8 |  |

**Table 2:** Clinicopathological features of studied patients

| (n=34) | SP+(23) | SP-(11) | Total (34) |
| --- | --- | --- | --- |
| **Age(years)** |  |  |  |
| >60 | 4 (17.39%) | 1 (9.09%) | 5 (14.7%) |
| <60 | 19 (82.6%) | 10 (90.9%) | 29 (85.29%) |
| **Menopause status** |  |  |  |
| Pre | 12 (52.17%) | 7 (63.63%) | 19 (55.88%) |
| Post | 11 (47.82%) | 4 (36.36%) | 15 (44.11%) |
| **Tumor size(cm)** |  |  |  |
| <2 | 3 (13.04%) | 0 | 3 (8.82%) |
| 2-5 | 15 (65.2%) | 8 (72.7%) | 23 (67.64%) |
| >5 | 5 (21.73%) | 3 (27.2%) | 8 (23.52%) |
| **Grade** |  |  |  |
| I (well diff) | 2 (8.69%) | 1 (9.09%) | 3 (8.82%) |
| II (mod) | 7 (30.43%) | 1 (9.09%) | 8 (23.52%) |
| III (poor) | 14 (60.8%) | 9 (81.8%) | 23 (67.64%) |
| **TNM** |  |  |  |
| PT1 | 4 (17.39%) | 1 (9.09%) | 5 (14.7%) |
| PT2 | 15 (65.2%) | 8 (72.7%) | 23 (67.64%) |
| PT3 | 2 (8.69%) | 2 (18.2%) | 4 (11.76%) |
| PT4 | 2 (8.69%) | 0 | 2 (5.88%) |
| **Tumor type** |  |  |  |
| IDC | 15 (65.2%) | 5 (45.5%) | 20 (58.82%) |
| DCIS | 1 (4.34%) | 6 (54.5%) | 7 (20.58%) |
| ILC | 2 (8.69%) | 0 | 2 (5.88%) |
| IDC+DCIS | 5 (21.7%) | 0 | 5 (14.7%) |
| **ER status** |  |  |  |
| +ve | 19 (82.6%) | 9 (81.81%) | 28 (82.35%) |
| -ve | 4 (17.39%) | 2 (18.18%) | 6 (17.64%) |
| **PR status** |  |  |  |
| +ve | 19 (82.6%) | 9 (81.81%) | 28(82.35%) |
| -ve | 4 (17.39%) | 2 (18.18%) | 6 (17.64%) |
| **HER2/neu status** |  |  |  |
| +ve | 18 (78.26%) | 4 (36.4%) | 22 (64.7%) |
| -ve | 5 (21.7%) | 7 (63.4%) | 12 (35.29%) |
| **Ki-67 status** |  |  |  |
| +ve | 23 (100%) | 7 (63.4%) | 30 (88.23%) |
| -ve | 0 | 4 (36.4%) | 4 (11.76%) |
| **Distant metastasis** |  |  |  |
| present | 3 (13.04%) | 1 (9.09%) | 4 (11.76%) |
| absent | 18 (78.26%) | 9 (81.8%) | 27 (79.41%) |
| unknown | 2 (8.69%) | 1 (9.09%) | 3 (8.82%) |
| **Lymph node metastasis (axillary)** |  |  |  |
| 1-3 Lymph nodes | 2 (8.69%) | 2 (18.2%) | 4 (11.76%) |
| >4 Lymph nodes | 6 (26.08%) | 3 (27.3%) | 9 (26.47%) |
| **absent** | 15 (65.2%) | 6 (54.5%) | 21 (61.76%) |

## RESULTS AND DISCUSSION

### SP/NK-1R expression

Expression of SP and NK-1R was detected to be cytoplasmic. Expression of SP showed 68% (23) of the BC cases to be positive (table 3). Cases of well differentiated (WD) carcinoma had clear cells with cytoplasm and nucleus (Figure 1A) and most of them (66.6%) were SP positive (Figure 1C, table 3). In moderately differentiated (MD) cases little morphology of cells has been disrupted but so far they can be recognized (Figure 1D). In poorly differentiated (PD) cases (14 cases, 60.8%) maximum intensity (+3) of SP was observed (Figure 1G; table 2) whereas (7 cases, 87.5%) (Figure 1F, table 3) were MD with +2 intensity of SP expression and low intensity (+1) was seen in WD cases (2 cases) (Figure 1C, table 3). In poorly differentiated cases the cells morphology was extremely distorted, and cells couldn’t be simply distinguished (Figure 1I). Immunohistochemical staining for NK-1R was completed in a small number of core biopsies. The expression of NK-1R was similarly found to be related with the progression of BC. Its expression was high in MD and PD cases (Figure 1B,E,H).

**Figure 1:**
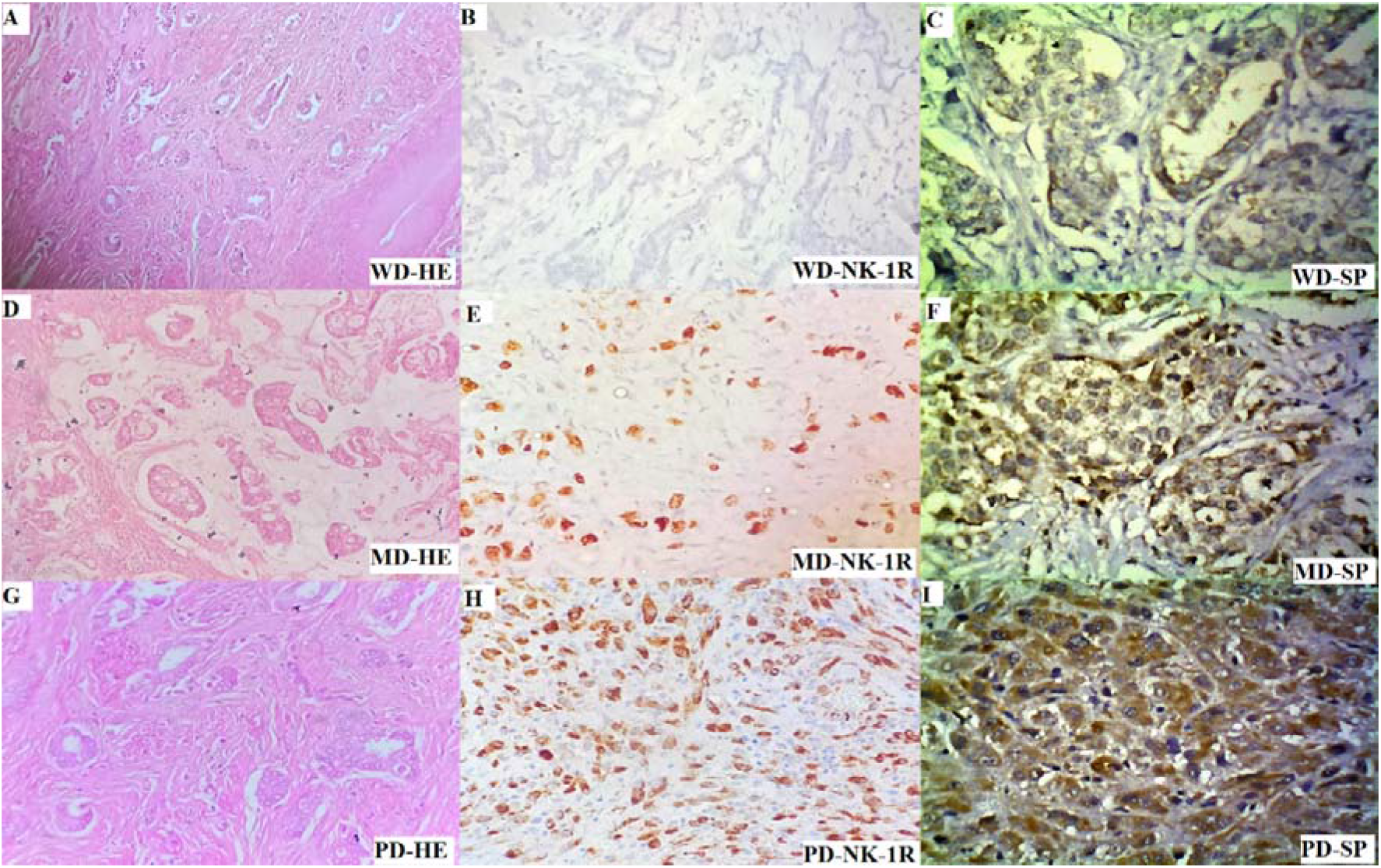
BC at 40X (A)WD-BC Hematoxylin Eosin staining (B) grade 1, NK-1R negative (C) SP weakly positive +1; (D) MD-BC Hematoxylin Eosin staining (E) MD, grade 2, NK-1R moderately positive, +2, 40% cells showing positive stain (F) MD, grade 2, SP moderately positive, +2; (G) PD-BC Hematoxylin Eosin staining (H) PD, grade 3, strongly SP positive, +3, 90% SP positive cells (I) PD, grade 3, strong positive, +3, 85% cells showing positive stain.

**Table 3:** Clinical Classification of breast cancer cases and its association with SP expression

| Types of breast cancer | SP+ | SP- |
| --- | --- | --- |
| Luminal A (ER/PR +, HER2 -) | 5 | 7 |
| Luminal B(ER/PR+, HER2 +) | 14 | 2 |
| ER/PR-, HER2 + | 4 | 2 |
| **Total cases** | 23 | 11 |

Clinicopathological patients

### Association of SP and patient characteristics with clinicopathological features of BC patients

Maximum number of the SP positive cases 19/23 (82.6%) belonged to the age group of < 60 years. 12/23 (52.17%) SP positive cases belonged to premenopausal females and 11/23 (47.82%) from postmenopausal females. Most cases, 15/23 (65.2%) had tumor sizes ranged between 2-5 cm. 14/23 (60.8%) cases of PD or grade III (group A), 7/23 (30.43%) cases of MD or grade II (group B) and 2/23 (8.69%) cases of WD-BC or grade I (group C) were SP positive. According to the TNM staging, 15/23 (65.2%) SP positive cases had PT2 stage. According to tumor type, 15/23 (65.2%) SP positive cases were invasive ductal carcinoma. Distant metastatis was absent in majority (18/23, 78.26%) of the SP positive cases. Axillary lymph node metastasis was also absent in (15/23, 65.2%) cases (table 3).

#### 1.1. Distribution of positive cases of SP according to the BC classification

Interpreting from the division of BC, 5/23 (21.73%) SP positive cases belonged to Luminal A group (ER/PR+, HER2-), 14/23 (60.8%) cases belonged to Luminal B (ER/PR+, HER2+) group and 4/23 (17.39%) to (ER/PR-, HER2+) group of BC (table 4).

**Table 4:** Expression and scoring of ER, PR and HER-2 in SP negative breast cancer cases

| SP Negative cases (n=11) |  |  |  |  |  |  |  |  |  |  |  |  |  |
| --- | --- | --- | --- | --- | --- | --- | --- | --- | --- | --- | --- | --- | --- |
| Age | Grade | Histo-opinion | Expression | %of cell stain | Intensity of stain | ALLRED SCORE | Expressi on | %of cell stain | Intensity of stain | ALLRE D SCORE | Expressiom | TNM | Size |
|  |  |  | ER status |  |  |  | PR status |  |  |  | HER2 status |  |  |
| 50 | 1 | DCIS-CB | +++ | 5 | 3 | 8 | ++ | 4 | 3 | 7 | - | PT1 | >5 |
| 40 | 3 | IDCIS | - | - | - | - | - | - | - | - | + | PT3 | >5 |
| 40 | 3 | IDCIS | + | - | - | - | + | - | - | - | - | PT2 | 2-5 |
| 37 | 3 | IDCIS | + | 2 | 2 | 4 | ++ | 3 | 3 | 6 | +++ | PT2 | 2-5 |
| 42 | 3 | IDC | +++ | 5 | 3 | 8 | + | 2 | 2 | 4 | +++ | PT2 | 2-5 |
| 47 | 3 | IDC | ++ | 3 | 3 | 6 | ++ | 4 | 3 | 7 | - | PT2 | 2-5 |
| 35 | 2 | DCIS(nipple involve) | - | - | - | - | - | - | - | - | + | PT2 | 2-5 |
| 40 | 3 | IDCIS | + | - | - | - | + | - | - | - | - | PT2 | 2-5 |
| 57 | 3 | IDC | ++ | 2 | 2 | 4 | + | 6 | - | - | - | PT2 | 2-5 |
| 72 | 3 | IDC | ++ | 4 | 2 | 6 | + | - | - | - | - | PT3 | >5 |
| 57 | 3 | IDC | +++ | 5 | 3 | 8 | ++ | 4 | 3 | 7 | - | PT2 | 2-5 |

#### 1.2. SP association with ER, PR, HER2 and Ki-67

ER was SP positive in 19/23 (82.6%); PR was positive in 17/23 (73.9%), HER-2 was positive in 18/23 (78.2%) of SP positive cases (Figure 2A-F). Ki-67 was positive in all the cases (Figure 2G,H) (table 3, 5 and 6). H scoring, ALLRED scoring and expressions of SP, ER, PR, HER2/neu and intensities of all stains are all mentioned in table 5,6.

**Figure 2:**
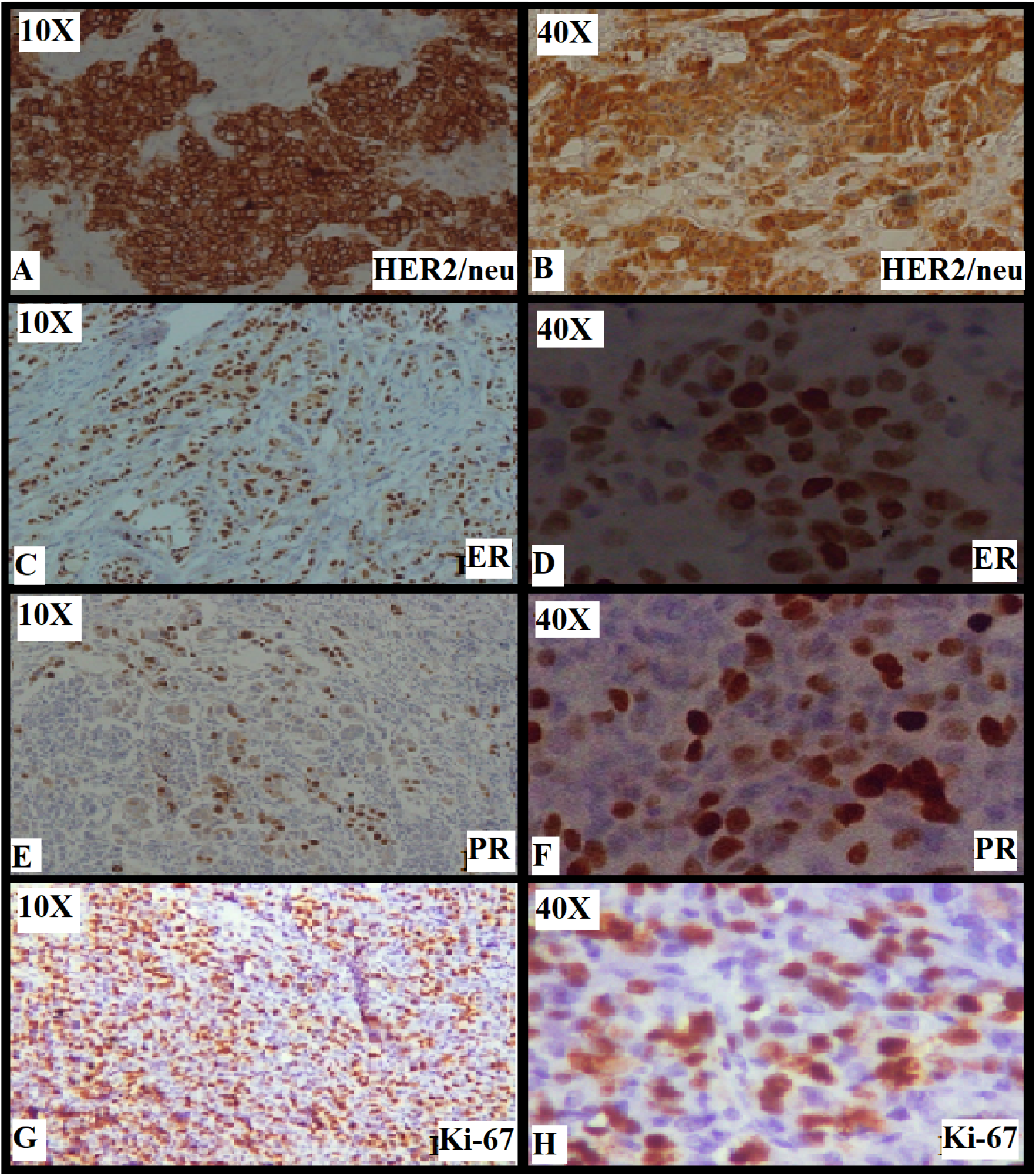
Staining with routine diagnostic markers for **BC (A,B)** HER-2 strongly positive, 10X and 40X; **(C,D)** ER strongly positive, 10X and 40X; **(E,F)** PR strongly positive, 10X and 40X; **(G,H)** Ki-67 proliferative marker, strongly positive, 10X and 40X

**Table 5:** Expression and scoring of ER, PR and HER2 in SP positive breast cancer cases

| SP positive cases (n=23) |  |  |  |  |  |  |  |  |  |  |  |  |  |  |  |
| --- | --- | --- | --- | --- | --- | --- | --- | --- | --- | --- | --- | --- | --- | --- | --- |
| Age | Grade | Histo-opinion | SP Expression | H score | Expression | % of cell stain | Intensity of stain | ALLRED SCORE | Expression | % of cell stain | Intensity of stain | ALLRED SCORE | Expression | TNM | Size(cm) |
|  |  |  |  |  | ER status |  |  |  | PR status |  |  |  | HER2 status |  |  |
| 33 | 2 | IDC | +++ | 140 | - | - | - | - | - | - | - | - | ++ | PT2 | <2 |
| 58 | 3 | IDC=DCIS | +++ | 190 | +++ | 5 | 3 | 8 | +++ | 5 | 3 | 8 | +++ | PT4 | 2-5 |
| 57 | 3 | IDC | +++ | 240 | +++ | 5 | 3 | 8 | +++ | 5 | 3 | 8 | +++ | PT2 | 2-5 |
| 34 | 3 | IDC | +++ | 240 | + | 2 | 2 | 4 | ++ | 4 | 2 | 6 | - | PT2 | <2 |
| 50 | 3 | IDC+DCIS | +++ | 150 | +++ | 5 | 3 | 8 | ++ | 4 | 2 | 6 | ++ | PT4b | 2-5 |
| 62 | 3 | IDC+DCIS | ++ | 120 | + | 2 | 3 | 5 | ++ | - | - | 6 | ++ | PT2 | 2-5 |
| 54 | 3 | IDC | +++ | 210 | +++ | 5 | 3 | 8 | +++ | 5 | 3 | 8 | +++ | PT2 | 2-5 |
| 33 | 2 | IDC | ++ | 160 | +++ | 5 | 3 | 8 | ++ | 4 | 3 | 7 | ++ | PT2 | 2-5 |
| 37 | 3 | IDC+DCIS | +++ | 210 | ++ | 4 | 2 | 6 | +++ | 5 | 3 | 8 | +++ | PT1c | >5 |
| 67 | 2 | IDC+E-DCIS | + | 140 | + | 3 | 1 | 4 | - | + | - | - | - | PT3 | 2-5 |
| 63 | 1 | DCIS | + | 60 | + | 2 | 2 | 4 | ++ | 3 | 3 | 6 | - | PT1 | >5 |
| 32 | 3 | IDC | + | 180 | +++ | 5 | 3 | 8 | +++ | 5 | 3 | 8 | +++ | PT2 | <2 |
| 33 | 3 | IDC | + | 70 | + | 3 | 2 | 5 | ++ | 3 | 3 | 6 | - | PT2 | 2-5 |
| 42 | 2 | ILC | + | 150 | +++ | 5 | 3 | 8 | ++ | 4 | 3 | 7 | ++ | PT2 | >5 |
| 57 | 3 | IDC | ++ | 140 | - | - | - | - | - | - | - | - | ++ | PT2 | 2-5 |
| 37 | 3 | IDC | ++ | 150 | - | - | - | - | - | - | - | - | + | PT2 | 2-5 |
| 47 | 2 | IDC | ++ | 180 | + | 3 | 1 | 4 | - | + | - | 3 | + | PT2 | 2-5 |
| 34 | 3 | IDC | +++ | 160 | + | 2 | 2 | 4 | + | 3 | 1 | 4 | - | PT1 | >5 |
| 50 | 3 | IDC | +++ | 210 | + | 2 | 2 | 4 | ++ | 5 | 2 | 7 | ++ | PT3 | 2-5 |
| 67 | 1 | IDC | + | 160 | +++ | 5 | 3 | 8 | ++ | 4 | 3 | 7 | ++ | PT2 | 2-5 |
| 42 | 3 | IDC | +++ | 240 | +++ | 5 | 3 | 8 | ++ | 4 | 3 | 7 | ++ | PT1c | >5 |
| 32 | 2 | ILC | ++ | 180 | +++ | 5 | 3 | 8 | +++ | 5 | 3 | 8 | ++ | PT2 | 2-5 |
| 50 | 2 | IDC | ++ | 160 | - | - | - | - | - | - | - | - | + | PT2 | 2-5 |

For the first time it is demonstrated that SP is not only overexpressed, but it is also involved in the progression of BC. It is found to be associated with poor prognosis and advancement of disease as reported by a previous study [19]. BC cells may release SP after binding to its receptor, NK-1R, as a possible mechanism, it may lead to proliferation [19], migration [21] and angiogenesis [25]. SP may also cause inflammation by enhancing the permeability of brain blood barrier (BBB) [26]. Subsequently, BC cells migrate and metastasize.

Similar findings were observed in our study except that we evaluated SP and NK-1R both in tissue but in previous study[27], only NK-1R was evaluated in tissue. There is little contradiction in SP evaluation: in our study we observed an increased expression with increasing grade of tumor while in previous study, no difference among the grades was observed but previous was only performed on serum. we revealed the SP expression in all grades of breast cancer which was commonly positive and the intensity increased with advancing grade. It demonstrates that SP expression is associated with the poor prognosis and aggression of this illness. Our outcomes are in concordance with the earlier studies on BC, which showed SP overexpression [19]. SP discharge from BC cells in response to nociceptive stimuli, whose consequences result in proliferation [19], metastasis and vasculogenesis [21] by functioning of autocrine and origins inflammation by paracrine role. SP rises the absorptivity of blood Brain Barrier (BBB) [25, 26]. Advanced grade of BC showed higher intensity of SP expression, they can be involved in metastasis.

When more SP is released, it can decrease the apoptosis subsequently [28] by modulating the immune markers IL4, IL6 and IL10 [29], resulting in unrestrained cell division, cell progression and prominent to cancer metastasis. All these mechanisms are carried by increased cellularity in human tenocytes [30] resulting in binding of SP to NK-1R. SP has been described to phosphorylate the Akt, antiapoptotic protein kinase [31]. SP has been studied in bone marrow stem cells showing proliferative effects [32] but it has to be considered in detail factor in cancer.

Previously, we had demonstrated the immunohistochemical expression of SP in the sudden fetal and infant deaths and neuropathology [33–36]. We also established SP expression in Oral Squamous Cell Carcinoma (OSCC), where SP strong expression was found to be related with the progression of OSCC and aided as a diagnostic marker [22]. It was directly related to the grade of cancer i.e. intensity of expression increased with the increasing grade. An *insilico* analysis by us also revealed the possible involvement of Tachykinin 1 (Tac1) gene, a gene for SP, in cancer [37]. In another study, SP/NK-1R system is found to be associated with colorectal cancer progression and prognosis [38].

Tachykinin family is the largest peptide family, its members bind to G-protein coupled receptors at the cells of destination. Hence, a signaling cascade is initiated, leading to mitogen activated protein kinase activation, mobilization of calcium and phosphoinsitide hydrolysis. Tumor microenvironment plays a crucial role in this regard and SP carries its role by binding to NK-1R[25]. SP is found to be important for the viability of cancer cells and NK-1R has been observed to be more expressed in these cells [39]. SP and NK-1R expresses more with the progression of several diseases[18, 40]. Our study is in accordance with these studies and we observed an overexpression of SP in grade III and intermediate expression in grade II.

Overexpression of SP and NK-1R was also observed in pre-cancerous epithelium and it was proposed that it has contribution towards early carcinogenesis by increasing cell growth, cell division [41], however, in current study, this trend was found in later stage of disease. NK-1R antagonists may inhibit cellular growth, proliferation and metastasis. It may have therapeutic role for cancer treatment by inhibiting neo-angiogenesis and vascularization. It may be explored for potential as antitumor drugs [18]. It may block the signal transduction network in cancer microenvironment and reduce the proliferation of tumor cells[40]. By contrast, NK-1R antagonists concentration dependent manner counteract SP pathophysiological functions. So, NK-1R antagonists may inhibit BC cellular growth, proliferation [19] and migration (for invasion and metastasis) [21]. It may have therapeutic role for cancer treatment by inhibiting neoangiogenesis and vascularization. It may be explored for potential as antitumor drugs [18]. It may block the signal transduction network in cancer microenvironment and reduce the proliferation of tumor cells [40].

In a recent case report published by Munoz M, a patient with chronic obstructive pulmonary disease and non-small cell lung carcinoma was treated with radiotherapy plus NK-1R antoagonist, aprepitant, for 45 days. The patient remained in good health, with no side effects and the tumor volume also decreased[42]. Further research and clinical trials must be carried out in order to fully reveal the beneficial effects of NK-1R antagonists in the treatment of patients suffering from BC. NK-1R antagonists can help in inhibition of various cancers by blocking angiogenesis[43].

## Conclusion

We hereby conclude that increased intensity and overexpression of Substance P and NK-1R is associated with poor prognosis in BC. SP/ NK-1R may also be explored further as a potential diagnostic biomarker for BC to differentiate the grades.

## Authors’ contributions

**RM** designed and planned the study, wrote the main manuscript and supervised the project

**SAG** critically reviewed the manuscript and facilitated this research

**AH** collected samples, medical history and did data analysis

**IT** gave histopathology opinion for the samples and scoring

**RUK** gave histopathology opinion for the samples and scoring

**MEB** edited and approved the final manuscript

**JA** critical review

**SS** Collected the samples, processed, stained and contributed in write up

**FJA** analysed the work, edited and approved the final manuscript

**MM** gave expert opinion in conducting the experimental work

All the authors have read the manuscript and finally approved

